# Hypothalamic perineuronal nets are regulated by sex and dietary interventions

**DOI:** 10.1101/2021.03.07.434286

**Authors:** Nan Zhang, Zili Yan, Hailan Liu, Meng Yu, Yang He, Hesong Liu, Chen Liang, Longlong Tu, Lina Wang, Na Yin, Junying Han, Nikolas Scarcelli, Yongjie Yang, Chunmei Wang, Tianshu Zeng, Lu-Lu Chen, Yong Xu

**Affiliations:** Children’s Nutrition Research Center, Department of Pediatrics, Baylor College of Medicine, Houston, TX, 77030; Department of Endocrinology, Union Hospital, Tongji Medical College, Huazhong University of Science and Technology, Wuhan, China; Hubei Provincial Clinical Research Center for Diabetes and Metabolic Disorder, Wuhan, China; Department of Molecular and Cellular Biology, Baylor College of Medicine, Houston, TX, 77030

**Author notes:** To whom correspondence should be addressed: Yong Xu, PhD, MD; 1100 Bates Street #8066, Houston, TX 77030, USA, Mail stop code: MCB320. LEAD AUTHOR: YONG XU.

## Abstract

Perineuronal nets (PNNs) are widely present in the hypothalamus, and are thought to provide physical protection and ion buffering for neurons, and regulate their synaptic plasticity and intracellular signaling. Recent evidence indicates that PNNs in the mediobasal hypothalamus plays an important role in the regulation of glucose homeostasis. However, whether and how hypothalamic PNNs are regulated are not fully understood. In the present study, we examined whether PNNs in various hypothalamic regions in mice can be regulated by sex, gonadal hormones, dietary interventions, or their interactions. We demonstrated that gonadal hormones are required to maintain normal PNNs in the arcuate nucleus of hypothalamus in both male and female mice. In addition, PNNs in the terete hypothalamic nucleus display a sexual dimorphism with females higher than males, and high-fat diet feeding increases terete PNNs only in female mice but not in male mice. On the other hand, PNNs in other hypothalamic regions are not influenced by sex, gonadal hormones or dietary interventions. In summary, we demonstrated that hypothalamic PNNs are regulated in a region-specific manner and these results provide a framework to further investigate the potential functions of PNNs in regulating energy/glucose homeostasis at the interplay of sex, gonadal hormones and diets.

## Introduction

Obesity is now recognized as a serious global health problem due to its increasing prevalence and comorbidities, e.g. the metabolic syndrome. The WHO reported that over 650 million adults worldwide were obese in 2016 and 40 million children under the age of 5 were overweight or obese in 2018. In U.S., the prevalence of adult obesity was 42.4% in 2017~2018 according to the Centers for Disease Control and Prevention. The etiology of human obesity is not fully understood, and effective treatments for obesity and associated metabolic disorders are limited. Recent studies revealed genetic and epigenetic basis for variations in human body mass index (BMI) (1–3), and strikingly the majority of BMI-associated genetic variants affect genes that are enriched in the brain (3,4). In particular, the brain hypothalamus receives metabolic and/or hormonal signals reflecting the body’s nutritional status and coordinates neuroendocrine and behavioral responses to maintain body weight balance (5–7). Indeed, various genetic variants associated with human obesity have been demonstrated to cause energy and/or glucose dysregulations through impairing functions of neurons and/or neurocircuits within the hypothalamus (8).

Perineuronal nets (PNNs) in the brain are condensed glycosaminoglycan-rich extracellular matrix structures (9). PNNs typically enmesh neurons in defined circuits, and are thought to provide physical protection and ion buffering for neurons, and regulate their synaptic plasticity and intracellular signaling (9). PNNs abundantly exist in the forebrain regions, e.g. the cortex and the hippocampus, and the PNNs levels in these regions can be regulated by animals’ experience, and have been implicated in various neurobiological disorders such as schizophrenia, bipolar disorder, Alzheimer’s disease and addictions. Recent evidence indicates that chemical disruption of PNNs in the mediobasal hypothalamus significantly blunts the glucose-lowering effects of central action of fibroblast growth factor 1 (FGF1) in obese Zucker diabetic fatty (ZDF) rats (10). Thus, hypothalamic PNNs may also play important roles in the regulation of energy and/or glucose homeostasis. Indeed, abundant PNNs are present in multiple hypothalamic regions (11,12), but whether and how these hypothalamic PNNs are regulated are not fully understood.

In the present study, we sought to examine whether PNNs in various hypothalamic regions can be regulated by dietary interventions in mice. Since many hypothalamic regions or neural populations are sexually dimorphic and/or regulated by gonadal hormones (13,14), we also explored the potential effects of sex and/or gonadal hormones on PNN levels. Our results revealed that sex, gonadal hormones and/or dietary interventions can regulate PNNs in a region-specific manner and provide a framework to further investigate the potential functions of PNNs in regulating energy/glucose homeostasis at the interplay of sex, gonadal hormones and diets.

## METHODS

### Study Animals

C57BL6j male and female mice were group-housed respectively until 14 weeks of age in a temperature-controlled room under a 12h:12h light-dark cycle with ad lib access to regular chow diet (5V5R-Advanced Protocol PicoLab Select Rodent 50 IF/6F, PicoLab) and water. Forty nine mice are divided into 16 groups by four factors (1) sex: male or female groups; (2) surgeries: receiving sham or castration (CAST)/ovariectomy (OVX) surgeries at 14 weeks of age (see below); (3) dietary interventions: chronic feeding with chow or a high-fat diet (HFD, D12492i Rodent Diet With 60 kcal% Fat, Research Diets, Inc.), (4) nutritional status: fed or fasted at the time of perfuse (see below). Care of all animals and procedures were approved by the Baylor College of Medicine Institutional Animal Care and Use Committee.

### Surgery

At the age of 14 weeks, either bilateral gonadectomy (CAST for males and OVX for females) or sham surgeries were executed in these mice. The mice were anesthetized with inhaled 2% isoflurane. For OVX in females, the dorsolateral incisions (2 cm) were made, followed by blunted separation of the subcutaneous fat, and incision (1.5cm) the abdominal wall to expose the reproductive tract. The ovaries were isolated and excised. The reproductive tract was returned to the peritoneum and the incisions on the abdominal wall were sutured using sterile Vicryl thread (size 6); the skin incisions were sutured by sterile Prolene thread. For sham surgeries in female mice, the procedures were the same, except that the ovaries were kept intact. For castration in males, a central vertical incision (3 cm) was made, followed by blunted separation of any subcutaneous fat to locate the vas deferens. The vas deferens were ligated with Vicryl; the testes were cut and taken away from the fat pad. The skin incisions were sutured by sterile Prolene thread. For sham surgeries in male mice, the procedures were the same, except that the testes were kept intact.

### Food intake, body weight, glucose and body composition

One week after the surgeries, mice were randomly divided into two groups to be either regular chow or HFD described above ad libitum for four weeks. Food intake, body weight and fed glucose were measured before the surgeries and once each week after the surgery for five weeks. The glucose was measured using the glucometer via the tail vein; to avoid confounding effects from circadian clock and feeding behavior on the glucose level, we always measured blood glucose in the morning of the day and after a 2-hour short fasting to ensure the empty stomach. Body composition (fat mass and lean mass) was examined with the Bruker minispec mq10 MRS system before the surgeries and 5 weeks afterwards.

### Perfusion and WFA staining

Five weeks after the surgeries, mice were further divided into fasted and ad libitum groups before the perfusion. For the fasted group, mice were fasted overnight and then perfused in the next morning; for the ad libitum group, both food and water were provided ad libitum at all time before the perfusion. For the perfusion, mice were anaesthetized with inhaled isoflurane and perfused with saline followed by 10% formalin. The brain sections were cut at 25 μm and collected into five consecutive series. One series IS used for the following staining and quantification. Free-floating tissue sections were incubated overnight at room temperature for 22 hours using a 1:500 dilution of biotin-labelled Wisteria floribunda agglutinin (WFA) (L1516, Sigma-Aldrich) in PBS + 0.25% Triton X-100. The sections were then washed and incubated for 2 hours at room temperature in 1:500 Streptavidin protein 488 (21832, Invitrogen). Images were captured using a fluorescence microscope with 1s exposure time and no change of contrast for all pictures to visualize various hypothalamic nuclei, including the arcuate hypothalamic nucleus (ARH), the terete hypothalamic nucleus (TE), the paraventricular hypothalamic nucleus (PVH), lateral hypothalamus (LH), the anterior hypothalamus (AH), the ventromedial hypothalamic nucleus (VMH), the dorsomedial hypothalamic nucleus (DMH), and the perifornical area of the anterior hypothalamus (PeFAH). Fluorescence intensity of PNNs for each image was quantified using an established method by the macro plugin “Perineuronal net Intensity Program for the Standardization and Quantification of ECM Analysis” (PIPSQUEAK AI v5.3.9, Rewire Neuro, Inc.) (15).

### Statistics

Statistical analysis was done using SPSS (SPSS 22.0.0.0, IBM SPSS Statistics) and GraphPad Prism (GraphPad Prism 8.4.2, Graphpad software, LLC). Multivariate linear regression in SPSS was used to investigate the influence of sex, GDX and chronic dietary intervention on various metabolic parameters (cumulative food intake, body weight change, blood glucose change, fat change and lean change). If sex or GDX was found to have a significant effect on any parameter, the p value for the interaction of sex and GDX was tested. If any two factors were found to have significant effects, the p value for the interaction of these two factors was tested.

For the PNN fluorescence intensity, in addition to the sex, GDX and chronic diet intervention, we introduced one more factor (nutritional status) as mice were perfused either after an overnight fasting or ad libitum. Similarly, we first used the multivariate linear regression analyses in SPSS to examine effects of these 4 factors on PNNs fluorescence intensity in each hypothalamic region. If sex or GDX was found to have a significant effect on any parameter, we then used the two-way ANOVA to examine the interaction of sex and GDX. In case that sex had a significant effect, we used the GraphPad Prism to further examine effects of dietary intervention and nutritional status in each sex separately. P < 0.05 was considered statistically significant.

### Study approval

Care of all animals and procedures were approved by the Baylor College of Medicine Institutional Animal Care and Use Committee.

## Results

### Distribution of WFA-labelled PNNs in the mouse hypothalamus

We first surveyed the distribution of PNNs in the mouse hypothalamus using WFA immunostaining. Abundant WFA-labelled PNNs were detected in multiple hypothalamic regions, including the LH, the ARH, the VMH and the TE (Figure 1). In addition, we also observed modest WFA immunoreactivity in the AH, the PVH and the DMH (Figure 1). Consistent with a previous report (12), WFA-labelled PNNs were also observed in a newly identified hypothalamic area, namely the PeFAH (Figure 1A).

**Figure 1.**
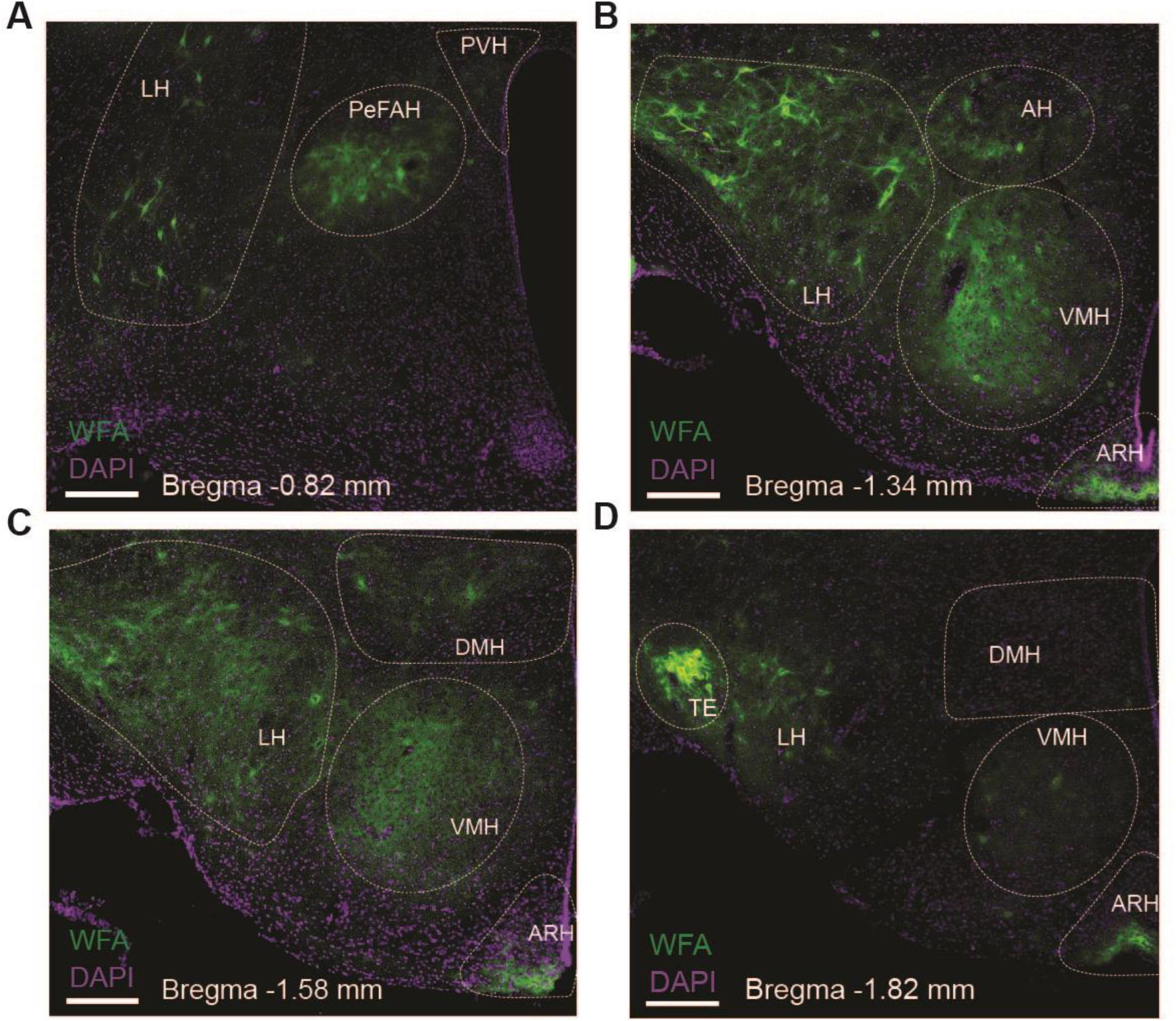
Distribution of WFA-labelled PNNs in mouse hypothalamus. Representative fluorescent microscopic images showing WFA-labelled PNNs (green) and DAPI counterstaining (purple) in a series of coronal brain sections at the level of −0.82 mm (A), −1.34 mm (B), −1.58 mm (C) and −1.82 mm (D) relative to the Bregma. Scale bars = 200 μm. AH, the anterior hypothalamus; ARH, the arcuate hypothalamic nucleus; DMH, the dorsomedial hypothalamic nucleus; LH, the lateral hypothalamus; PeFAH, the perifornical area of the anterior hypothalamus (PeFAH); PVH, the paraventricular hypothalamic nucleus; TE, the terete hypothalamic nucleus; VMH, the ventromedial hypothalamic nucleus.

### Effects of sex, gonadal hormones and dietary interventions on metabolic parameters

Given the essential roles of the hypothalamus in regulating energy homeostasis, we sought to examine potential effects of dietary interventions on the level of PNNs in the hypothalamus. In order to explore potential influence by sex and/or gonadal hormones, we included both male and female mice with or without gonadectomy (GDX) in these analyses. Briefly, male and female C57Bl6j mice underwent sham or GDX (castration for males and ovariectomy for females) at the 14 weeks of age, followed by 5-week chronic feeding with either a regular chow diet or HFD. As expected, the multivariate linear regression analysis revealed that compared to chow-fed groups, HFD feeding had profound effects on multiple metabolic parameters, including significantly increasing calorie intake, blood glucose, body weight gain and fat mass gain (Table 1). Notably, fat mass gain was significantly smaller in females than in males (Table 1). We also noted that lean mass gain was significantly larger in females than in males, and that GDX significantly reduced lean mass gain; in addition, there was a significant interaction between sex and GDX on this parameter (Table 1). In summary, multiple metabolic parameters in mice were influenced by sex, gonadal hormones, the dietary interventions, and/or their interactions, as reported by others (16,17).

**Table 1.**
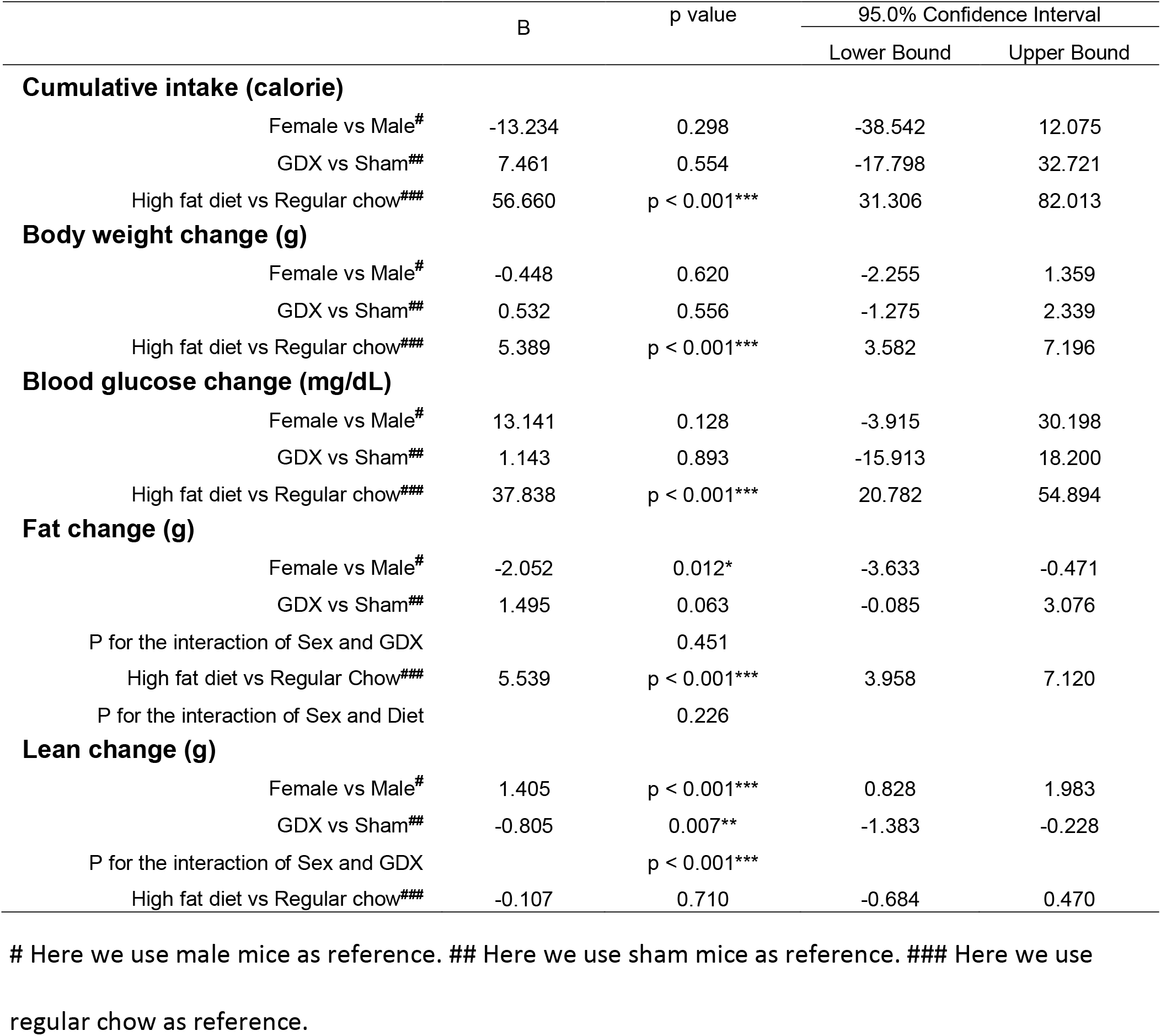
Effects of sex, GDX and diet on various metabolic parameters.

### Effects of gonadal hormones on PNN in the ARH

At the end of 5-week feeding, mice were then perfused either ad libitum or after an overnight fasting. We then used these fixed brain samples for WFA immunostaining and then quantified the level of WFA-labelled PNNs in each hypothalamic region in various conditions. In the ARH, the WFA fluorescence intensity was not significantly affected by chronic HFD feeding (vs. chow), or by the acute fasting (vs. ad libitum), as revealed by the multivariate linear regression analysis (Table 2). Interestingly, we noted that GDX significantly reduced WFA fluorescence intensity in the ARH compared to the sham group (Table 2 and Figure 2A–2E). A detailed analysis at different rostral-caudal levels revealed that GDX mainly reduced WFA in the rostral to medial ARH (−1.94 to −1.58 mm to Bregma) but the caudal ARH was not affected (Figure 2F). Because female and male mice experienced different GDX surgeries (OVX for females and CAST for males), we used a two-way ANOVA analysis to further analyze the interaction between sex and GDX. We found that the inhibitory effect of GDX was independent of sex; in other words, both OVX in females and CAST in males significantly reduced WFA fluorescence intensity in the ARH (Table 2 and Figure 2G). We also examined effects of dietary innervations on ARH PNNs in sham or in GDX groups, respectively, but did not detect any significant effect of by either chronic HFD feeding or by acute fasting (data not shown). Thus, these results indicate that endogenous gonadal hormones in both males and females are required to maintain normal PNNs in the ARH.

**Figure 2.**
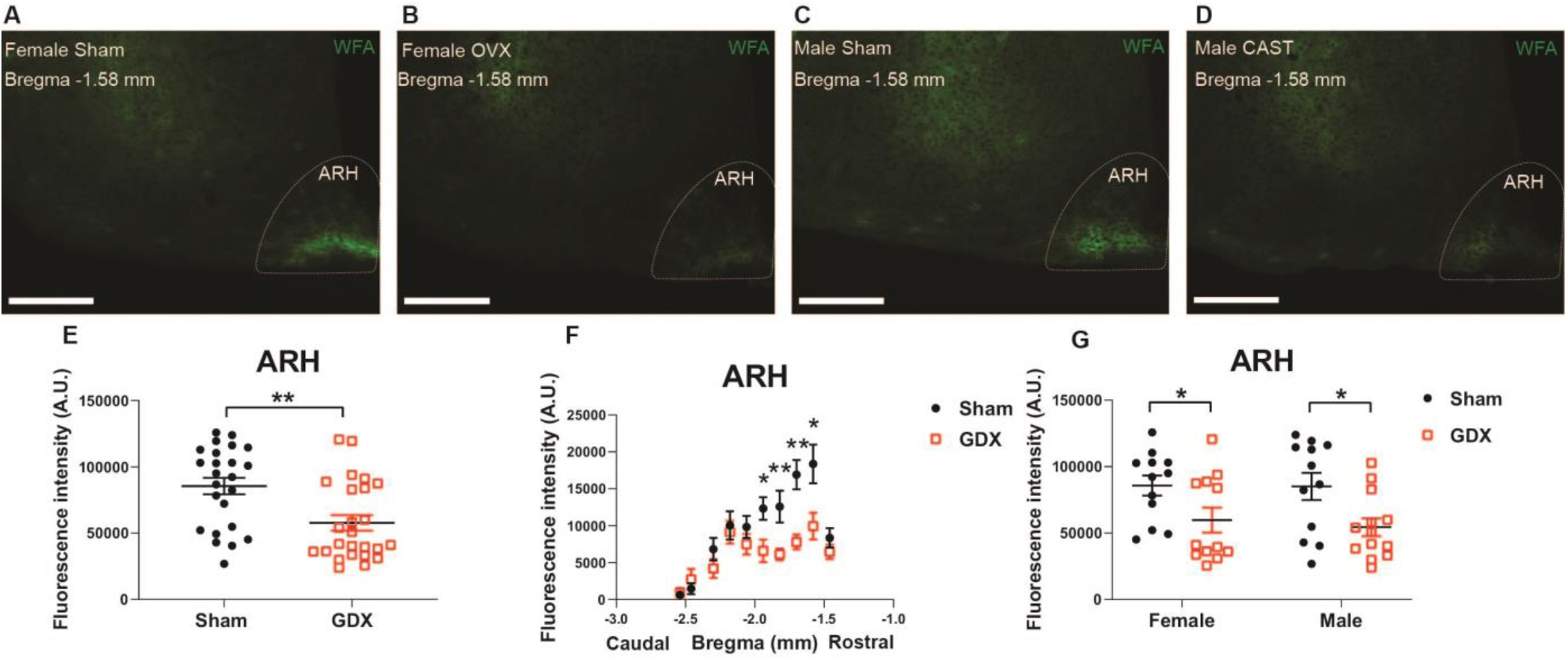
GDX reduces PNNs in the ARH. (A-D) Representative fluorescence microscopic images showing WFA-labelled PNNs (green) in the ARH of female sham (A), female OVX (B), male sham (C) and male CAST mice (D). Scale bars = 200 μm. ARH, the arcuate hypothalamic nucleus. (E) Quantification of the total WFA fluorescence intensity in the ARH from sham vs. GDX mice. N = 24 or 25 mice per group. Data are presented with mean ± SEM with individual data points. **, P<0.01 in unpaired two-tailed Student’s t-tests. (F) Quantification of the WFA fluorescence intensity at various rostral-caudal levels of the ARH from sham vs. GDX mice. N = 24 or 25 mice per group. Data are presented with mean ± SEM. *, P<0.05 and **, P<0.01 in unpaired two-tailed Student’s t-tests at each level. (G) Quantification of the total WFA fluorescence intensity in the ARH from female or male mice with sham or GDX surgeries. N = 12 or 13 mice per group. Data are presented with mean ± SEM with individual data points. *, P<0.05 in two-way ANOVA analysis followed by Holm-Sidak post hoc tests.

**Table 2.** Effects of sex, GDX, diet and fasting on PNNs in the ARH.

|  | B | p value | 95.0% Confidence Interval |  |
| --- | --- | --- | --- | --- |
|  |  |  | Lower Bound | Upper Bound |
| Female vs Male# | 3216.040 | 0.710 | -14091.082 | 20523.162 |
| GDX vs Sham## | -28525.814 | 0.002** | -45832.935 | -11218.692 |
| High fat diet vs Regular chow### | 5478.986 | 0.527 | -11858.261 | 22816.234 |
| Fast vs Ab libitum#### | 1070.635 | 0.902 | -16266.612 | 18407.883 |

### Effects of sex and HFD feeding on PNN in the TE

In the TE, female mice showed significantly higher WFA fluorescence intensity compared to males, as revealed by the multivariate linear regression analysis (Table 3 and Figure 3A–3E), and this sex difference existed in a rostral subdivision (−1.70 mm to Bregma) and a medial-caudal subdivision (−2.18 to −2.30 mm to Bregma) of the TE (Figure 3F). Using a two-way ANOVA analysis, we further found that the significant sex difference only existed in sham mice, but not in GDX mice (Figure 3G). Thus, these results indicate that gonad-intact female mice have higher PNNs in the TE than gonad-intact males.

**Figure 3.**
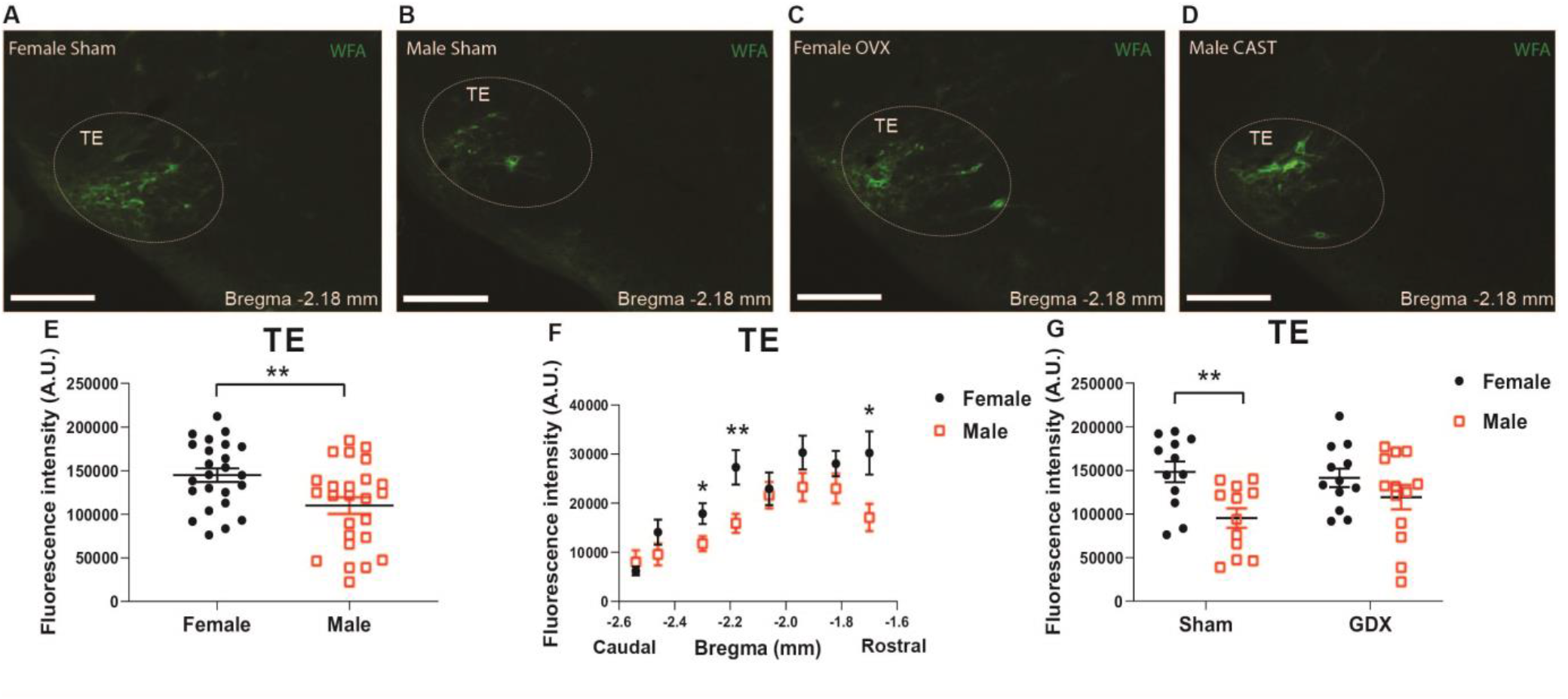
PNNs in the TE display a sexual dimorphism. (A-D) Representative fluorescence microscopic images showing WFA-labelled PNNs (green) in the TE of female sham (A), male sham (B), female OVX (C), and male CAST mice (D). Scale bars = 200 μm. TE, the terete hypothalamic nucleus. (E) Quantification of the total WFA fluorescence intensity in the TE from female vs. male mice. N = 24 or 25 mice per group. Data are presented with mean ± SEM with individual data points. **, P<0.01 in unpaired two-tailed Student’s t-tests. (F) Quantification of the WFA fluorescence intensity at various rostral-caudal levels of the TE from female vs. male mice. N = 24 or 25 mice per group. Data are presented with mean ± SEM. *, P<0.05 and **, P<0.01 in unpaired two-tailed Student’s t-tests at each level. (G) Quantification of the total WFA fluorescence intensity in the TE from female or male mice with sham or GDX surgeries. N = 12 or 13 mice per group. Data are presented with mean ± SEM with individual data points. **, P<0.01 in two way ANOVA analysis followed by Sidak post hoc tests.

**Table 3.** Effects of sex, GDX, diet and nutritional status on PNNs in the TE.

|  | B | p value | 95.0% Confidence Interval |  |
| --- | --- | --- | --- | --- |
|  |  |  | Lower Bound | Upper Bound |
| Female vs Male<sup>#</sup> | 37265.520 | 0.004** | 12278.584 | 62252.456 |
| GDX vs Sham<sup>##</sup> | 8878.365 | 0.478 | -16108.572 | 33865.301 |
| High fat diet vs Regular chow<sup>###</sup> | 3443.149 | 0.783 | -21587.281 | 28473.579 |
| Fast vs Ab libitum<sup>####</sup> | -2328.269 | 0.852 | -27358.698 | 22702.161 |

Given the clear influence by sex, we reanalyzed the effects of HFD feeding in each sex individually and used a two-way ANOVA analysis followed by Sidak tests to further explore potential interactions between chronic HFD feeding and acute fasting. In female mice fed ad libitum, chronic HFD feeding significantly enhanced WFA fluorescence intensity in the TE compared to chow-fed females, but this effect was not observed after an overnight fasting (Figure 4A–4E). This HFD-induced increases existed in a rostral-medial subdivision (−1.70 to −1.94 mm to Bregma) of the TE (Figure 4F). On the other hand, in male mice, neither chronic HFD feeding nor acute fasting induced any changes in WFA fluorescence intensity in the TE (Figure 4G). Thus, these results indicate that chronic HFD feeding can increase PNNs in the TE in female mice but not in male mice.

**Figure 4.**
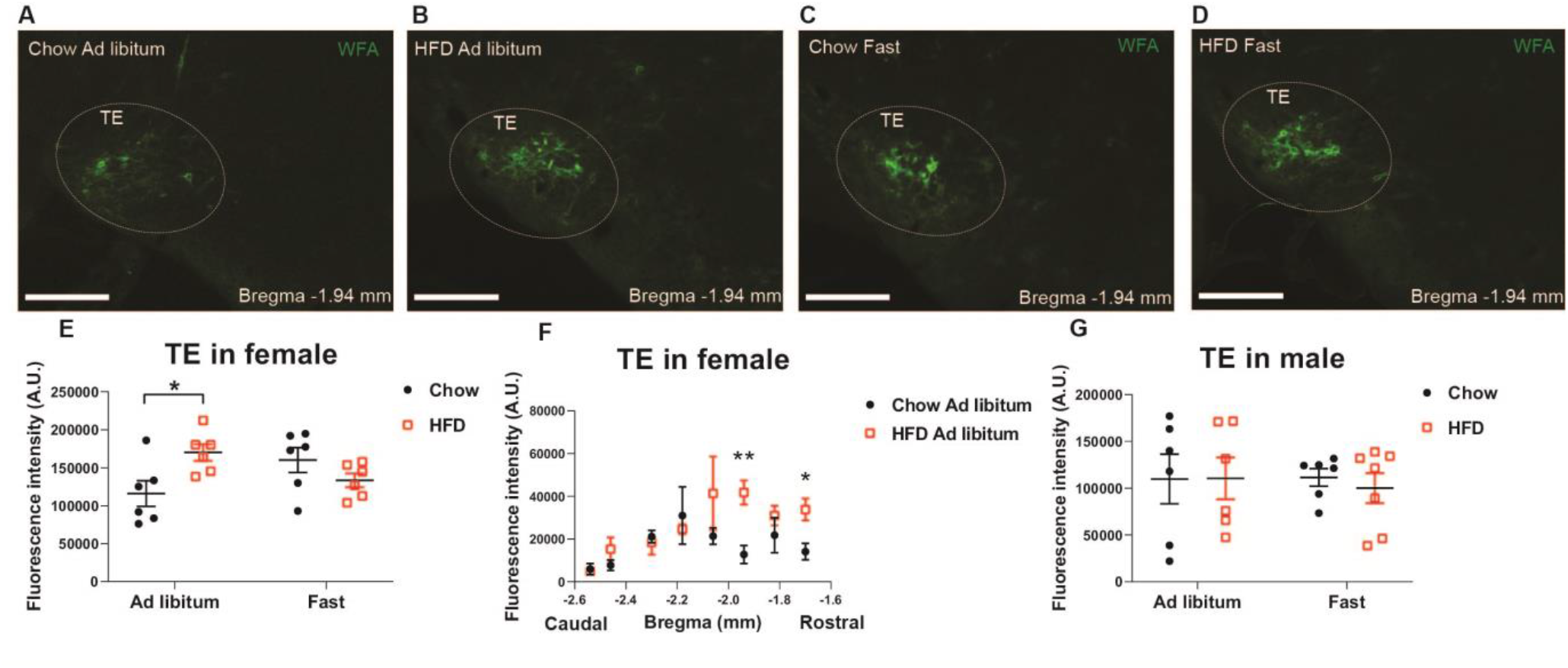
HFD feeding increases TE PNNs in female mice. (A-D) Representative fluorescence microscopic images showing WFA-labelled PNNs (green) in the TE of female mice that were fed chow ad libitum (A), HFD ad libitum sham (B), chow after an overnight fasting (C), and HFD after an overnight fasting (D). Scale bars = 200 μm. TE, the terete hypothalamic nucleus. (E) Quantification of the total WFA fluorescence intensity in the TE from female mice with various dietary interventions. N = 6 mice per group. Data are presented with mean ± SEM with individual data points. *, P<0.05 in two-way ANOVA analysis followed by Sidak post hoc tests. (F) Quantification of the WFA fluorescence intensity at various rostral-caudal levels of the TE from female mice fed with chow or HFD ad libitum. N = 6 mice per group. Data are presented with mean ± SEM. *, P<0.05 and **, P<0.01 in unpaired two-tailed Student’s t-tests at each level. (G) Quantification of the total WFA fluorescence intensity in the TE from male mice with various dietary interventions. N = 6 or 7 mice per group. Data are presented with mean ± SEM with individual data points. No significance was detected by in two way ANOVA analysis.

### Other hypothalamic regions

We also analyzed effects of sex, gonadal hormones, and/or dietary interventions on WFA fluorescence intensity in all other hypothalamic regions. The multivariate linear regression analysis revealed that WFA fluorescence intensity in the PVH, the PeFAH, the LH, the AH, the VMH or the DMH, was not significantly altered by these conditions (Table 4).

**Table 4.** Effects of sex, GDX, diet and nutritional status on PNNs in the PVH, PeFAH, LH, AH, VMH and DMH.

|  |  | B | p value | 95.0% Confidence Interval |  |
| --- | --- | --- | --- | --- | --- |
|  |  |  |  | Lower Bound | Upper Bound |
| PVH |  |  |  |  |  |
|  | Female vs Male<sup>#</sup> | 1093.487 | 0.785 | -6950.901 | 9137.874 |
|  | GDX vs Sham<sup>##</sup> | 1801.150 | 0.649 | -6123.285 | 9725.586 |
|  | High fat diet vs Regular chow<sup>###</sup> | 7111.050 | 0.082 | -948.109 | 15170.209 |
|  | Fast vs Ab libitum<sup>####</sup> | 1437.498 | 0.721 | -6621.661 | 9496.657 |
| PeFAH |  |  |  |  |  |
|  | Female vs Male<sup>#</sup> | 10127.575 | 0.447 | -16501.677 | 36756.827 |
|  | GDX vs Sham<sup>##</sup> | 12003.295 | 0.361 | -14228.883 | 38235.473 |
|  | High fat diet vs Regular chow<sup>###</sup> | 12968.496 | 0.332 | -13709.655 | 39646.646 |
|  | Fast vs Ab libitum<sup>####</sup> | 3879.818 | 0.771 | -22798.333 | 30557.969 |
| LH |  |  |  |  |  |
|  | Female vs Male<sup>#</sup> | 48723.960 | 0.118 | -12839.004 | 110286.924 |
|  | GDX vs Sham<sup>##</sup> | 25737.089 | 0.397 | -34907.897 | 86382.074 |
|  | High fat diet vs Regular chow<sup>###</sup> | -22043.519 | 0.475 | -83719.530 | 39632.491 |
|  | Fast vs Ab libitum<sup>####</sup> | -17950.520 | 0.560 | -79626.531 | 43725.490 |
| AH |  |  |  |  |  |
|  | Female vs Male<sup>#</sup> | 9830.509 | 0.347 | -11017.746 | 30678.764 |
|  | GDX vs Sham<sup>##</sup> | 6991.586 | 0.496 | -13545.796 | 27528.969 |
|  | High fat diet vs Regular chow<sup>###</sup> | 6079.521 | 0.560 | -14807.017 | 26966.059 |
|  | Fast vs Ab libitum<sup>####</sup> | 599.040 | 0.954 | -20287.498 | 21485.578 |
| VMH |  |  |  |  |  |
|  | Female vs Male<sup>#</sup> | 8833.935 | 0.639 | -28921.823 | 46589.693 |
|  | GDX vs Sham<sup>##</sup> | 6705.915 | 0.718 | -30486.859 | 43898.688 |
|  | High fat diet vs Regular chow<sup>###</sup> | -14866.204 | 0.432 | -52691.292 | 22958.884 |
|  | Fast vs Ab libitum<sup>####</sup> | -19095.426 | 0.314 | -56920.514 | 18729.662 |
| **DMH** |  |  |  |  |  |
|  | Female vs Male<sup>#</sup> | -2075.195 | 0.135 | -4826.375 | 675.985 |
|  | GDX vs Sham<sup>##</sup> | 267.179 | 0.843 | -2442.978 | 2977.336 |
|  | High fat diet vs Regular chow<sup>###</sup> | 1587.169 | 0.252 | -1169.064 | 4343.401 |
|  | Fast vs Ab libitum<sup>####</sup> | -487.298 | 0.723 | -3243.531 | 2268.934 |

## Discussions

One interesting observation of our study is that endogenous gonadal hormones are required to maintain normal PNNs in the ARH in both male and female mice. The ARH in the hypothalamus has been long believed to contain the first order neurons that respond to the peripheral signals, including leptin (18–20), insulin (21–23), ghrelin (24), asprosin (25) and estrogens (26–28). The ARH neurons include those expressing pro-opiomelanocortin (POMC) and those co-releasing agouti-related peptide (AgRP), neuropeptide Y (NPY), and GABA (referred to as AgRP neurons). POMC neurons synthesize and secret an anorexigenic peptide, α-melanocyte-stimulating hormone (α-MSH) which prevents overeating and body weight gain (29,30). On the other hand, AgRP neurons are orexigenic (31), and activation of AgRP neurons promotes eating even when mice are satiated (32,33). Recent evidence also indicates that non-AgRP GABAergic neurons in the ARH provide a redundant orexigenic mechanism to drive feeding and body weight gain (34). Notably, the majority of PNNs in the ARH enmeshes AgRP neurons and GABAergic neurons, with only a small portion enmeshing POMC neurons (11). Interestingly, reduced PNNs in the ARH is found in ob/ob mice deficient in the leptin gene and in ZDF rats with a mutated leptin receptor gene (10,11), highlighting an important role of leptin signaling in maintaining the normal PNNs in the ARH. Further, the central administration of FGF1 in ZDF rats can enhance PNNs in the ARH, associated with a prolonged glucose-lowering benefit (10). Importantly, chemical disruption of PNNs significantly shortens this glucose-lowering effect of FGF1 (10). Thus, PNNs in the ARH play an essential role in regulating glucose balance.

Here we demonstrated that PNNs in the ARH are significantly reduced by removal of the ovaries in female mice, suggesting that PNNs are also regulated by ovarian hormones. Supporting this notion, coadministration of both ovarian hormones (estrogen and progesterone), which simulates pregnancy, can trigger PNN formation in the medial preoptic area, a hypothalamic region normally devoid of PNNs in non-pregnant females (35). While the role of progesterone in energy and glucose homeostasis is not clear (14), estrogen is well established to provide many metabolic benefits, including suppressing food intake, increasing energy expenditure, reducing body weight gain, enhancing insulin sensitivity, and lowering blood glucose (16,36–41). Both estrogen receptor *α* (ERα) and estrogen receptor ß (ERß) are implicated in mediating these metabolic effects (42–44). Notably, abundant ERα is expressed by POMC neurons in the ARH (45–47), to a less extent by AgRP neurons (48,49), while ERβ expression in the ARH is minimal (50). We recently showed that intact ERα signals are required for the glucose-sensing functions of ARH neurons (51). Further, estrogen can enhance glutamatergic synapses onto POMC neurons (52), and increase their excitability (53,54). Importantly, we previously showed that selective deletion of ERα from POMC-lineage neurons in female mice results in hyperphagia and obesity (26), and attenuates estrogen-induced anorexia (55). Thus, these findings, together with our current observations that PNNs are reduced by ovariectomy, suggest that PNNs in the ARH may mediate, at least a portion of, estrogenic actions on energy and glucose homeostasis, and future investigations are warranted to test this possibility. Interestingly, we also observed similar reductions of PNNs in the ARH of male mice depleted with endogenous testosterone. Consistently, it has been reported that testosterone administration can stimulate PNN formation in the forebrain of song birds (56). However, it is worth noting that testosterone can be converted into estrogen by aromatase (57). Thus, it remains to be tested whether the testosterone-induced PNN expression is mediated by testosterone per se or by converted estrogen.

Another hypothalamic region where PNNs are regulated is the TE, where female mice showed higher PNNs than males. Importantly, this sex difference was only observed in gonad-intact mice, but was blunted in mice with gonads surgically removed. These results suggest that PNNs in the TE may be also regulated by gonadal hormones. Supporting this possibility, estrogen has been shown to increase oxytocin binding in the TE in rats (58), indicating potential actions of estrogen in this region. It has to be pointed out that, unlike other hypothalamic regions (e.g. the ARH, VMH, etc.), the TE has received very little attention with no more than 5 publications regarding this nucleus available in the PubMed. Early neuroanatomic studies reported that cell bodies of cholecystokinin neurons and calcitonin gene-related peptide (CGRP) neurons are found in the TE (59,60). In addition, the TE harbors nerve fibers that are immunoreactive for NPY, α-MSH, ß-endorphin, corticortropin-releasing hormone, galanin, and substance P (59,60), suggesting that the TE neurons receive a wide range of innervations and inputs from other neural populations. Interestingly, we found that in female mice only, PNNs in the TE was upregulated by chronic HFD feeding, while such effect was not observed after an overnight fasting. It is worth noting that among many hypothalamic regions expressing PNNs, the TE is the only region where PNNs can be regulated by dietary interventions. Considering that the TE receives inputs from NPY and α-MSH, both of which are implicated in the regulation of feeding and body weight, we suggest that TE neurons and the PNNs enmeshing these neurons may play an under-appreciated role in the context of energy homeostasis. This possibility warrants future investigations.

In summary, here we systemically assessed PNNs in multiple hypothalamic regions and explored potential regulations by sex, gonadal hormones, dietary interventions, and their interactions. We acknowledge that the limitation of the current study is its descriptive nature. However, we want to point out that our results filled a gap of knowledge in the relevant field and provided a framework to develop new hypotheses and design future studies to determine the physiological functions of hypothalamic PNNs and to explore the regulatory mechanisms. For example, the strong regulations by gonadal hormones on PNNs in the ARH suggest that gonadal hormones may act through PNNs in the ARH to regulate energy and/or glucose homeostasis. Further, given that PNNs in the ARH are required to mediates metabolic benefits of FGF1 (10), the efficacy of FGF1 may need to be examined in animals depleted of gonadal hormones, which would provide important pre-clinical data for potential applications of FGF1 or its analogs in aged men with testosterone deficiency or in post-menopausal women. In addition, the dietary regulations on PNNs in the TE suggest that this understudied hypothalamic nucleus may play a role in the regulation of feeding and body weight balance. Importantly, since these dietary regulations only existed in female mice but not in male mice, future studies need to include at least female animals, if not both. Only including male animals (a common practice in many laboratories) might miss potential discoveries about PNNs in the TE or TE neurons themselves.

## Author Contributions

NZ was involved in experimental design and most of procedures, data acquisition and analyses, and writing the manuscript. ZY assisted in data analysis. HL, MY, YH, HL, CL, LT, LW, NY, JH, NS, YY and CW assisted in in surgical procedures and production of study mice. LLC and TZ are involved in the study design and manuscript preparation. YX is the guarantors of this work and, as such, had full access to all the data in the study and take responsibility for the integrity of the data and the accuracy of the data analysis.

## Acknowledgements

The authors are grateful for the technical assistance provided by QW Zhang, SY Huang, DH Liu, B Yuan and YR Wang. The authors are grateful for the consultations provided by Dr. Michael Schwartz and Dr. Kimberly Alonge at University of Washington. The investigators were supported by grants from the NIH (K01DK119471 to CW), USDA/CRIS (51000-064-01S to YX) and the American Heart Association (20POST35120600 to YH).

## Data availability statement

All data generated or analyzed during this study are included in this published article.

